# A shape-constrained regression and wild bootstrap framework for reproducible drug synergy testing

**DOI:** 10.64898/2026.02.05.704019

**Authors:** Amir Asiaee, James P. Long, Samhita Pal, Heather H. Pua, Kevin R. Coombes

## Abstract

High-throughput drug combination screens require methods to identify synergistic pairs, yet widely used synergy scores lack statistical inference and can fail when parametric dose–response fits do not converge. We present SIR (Synergy via Isotonic Regression), a nonparametric framework that defines interaction as deviation from a monotone-additive null, fit by 2D isotonic regression. A degrees-of-freedom-corrected wild bootstrap yields calibrated p-values for each dose–response matrix. On DrugCombDB, SIR interaction surfaces achieve higher replicate concordance (median correlation 0.91 across 1,839 replicate pairs) than all baselines (0.53–0.74), while avoiding Loewe’s 20.9% and ZIP’s 3.6% failure rates. The fitted surface also predicts missing wells (median holdout RMSE 0.040). By replacing heuristic scores with calibrated effect sizes and p-values, SIR enables principled hit calling and error-rate control in large screens.

## 1 Introduction

Drug combinations are a cornerstone of cancer treatment, where monotherapy resistance and an ever-expanding pharmacopeia make rational combination selection critical. Combinations are routinely screened in vitro across dose matrices. Marginal dose-response curves (e.g., Hill or sigmoid models) are often fit to each drug individually to estimate potency and efficacy [1, 2]. These parametric fits are then used to define a “null” expected combination surface under various models of additivity, against which synergy is measured. However, marginal curve fitting can be sensitive to preprocessing and model misfit [3] and can fail to converge (produce no result) on real screening data [4].

Several null models have been proposed for quantifying synergy. Bliss independence [5] assumes that drugs affect cell survival probabilities independently and requires no curve fitting; HSA (highest single agent) simply compares the combination to the more active single agent at each dose pair; Loewe additivity [6, 7] and the Combination Index [8] assume dose equivalence and require fitted marginal curves; and ZIP [4, 9–11] assumes that drugs independently shift each other’s potency without changing curve shape, also requiring marginal fits. Each model encodes a different definition of “no interaction,” so the same dose-response matrix can be called synergistic by one model and antagonistic by another [12–16]. Moreover, all of these approaches produce pointwise scores at each dose pair, which are typically summarized by averaging across the matrix [4, 10]. While standard errors can in principle be computed at individual dose pairs, none of these methods provides a formal statistical test at the matrix level for whether overall interaction is present, nor uncertainty quantification for that conclusion. These limitations are increasingly consequential. Modern machine learning approaches often treat synergy scores as training labels [17–19], but if labels are inconsistent across models or unstable to noise and missingness, predictive models are incentivized to chase idiosyncrasies of the chosen scoring rule rather than reproducible biology. Recent community benchmarks have confirmed that the choice of synergy scoring model is a major source of variation in downstream predictions [19], underscoring the need for stable, statistically grounded alternatives. More fundamentally, without p-values or confidence intervals, it is impossible to distinguish a genuinely interacting combination from one whose synergy score is driven by measurement noise.

We introduce SIR (Synergy via Isotonic Regression), a framework that addresses these shortcomings. SIR replaces parametric curve fitting with monotone regression, a shape constraint that requires only that increasing dose does not decrease effect. Unlike Hill-curve fitting, monotone regression is a convex projection that always has a unique solution for any input data, so SIR never produces undefined outputs. SIR provides a calibrated p-value for each matrix by testing whether the dose-response surface deviates from a monotone-additive null, using a wild bootstrap [20] (a resampling method that generates pseudo-data by randomly flipping residual signs, preserving heteroscedasticity without parametric error assumptions). A global interaction statistic summarizes the entire matrix and, through bootstrap inference, supports reproducible discovery with error-rate control.

## 2 Results

### 2.1 Synergy labels show poor concordance across null models

To quantify disagreement among widely used baselines, we analyzed 391,652 dose– response matrices from DrugCombDB [21] for which all four baseline methods (Bliss, HSA, Loewe, ZIP) returned valid (non-NA) summary scores via SynergyFinder conventions [4]. Each score summarizes a full matrix by averaging pointwise synergy values across dose pairs. These matrix-level scores show strong dependence on the chosen null model: Bliss and ZIP correlate highly (Pearson *r* = 0.92), while Loewe correlates weakly with ZIP (*r* = 0.28) and Bliss (*r* = 0.33) (Fig. 1; pairwise scatterplots in Supplementary Fig. 4). Disagreement is not limited to scaling: for 21–34% of matrices, one method reports synergy (positive score) while the other reports antagonism (Fig. 1). Even “top hit” sets overlap modestly: among the top 5% most synergistic calls, the Loewe–ZIP Jaccard overlap is 0.36 (Fig. 1). These discrepancies imply that “synergy” labels used in downstream analyses, including ML training [17, 18], can vary substantially with the scoring model. The disagreement is not merely academic: a drug pair prioritized as a top hit by one method may rank in the bottom half by another, potentially leading to wasted experimental follow-up or missed therapeutic opportunities.

**Fig. 1.**
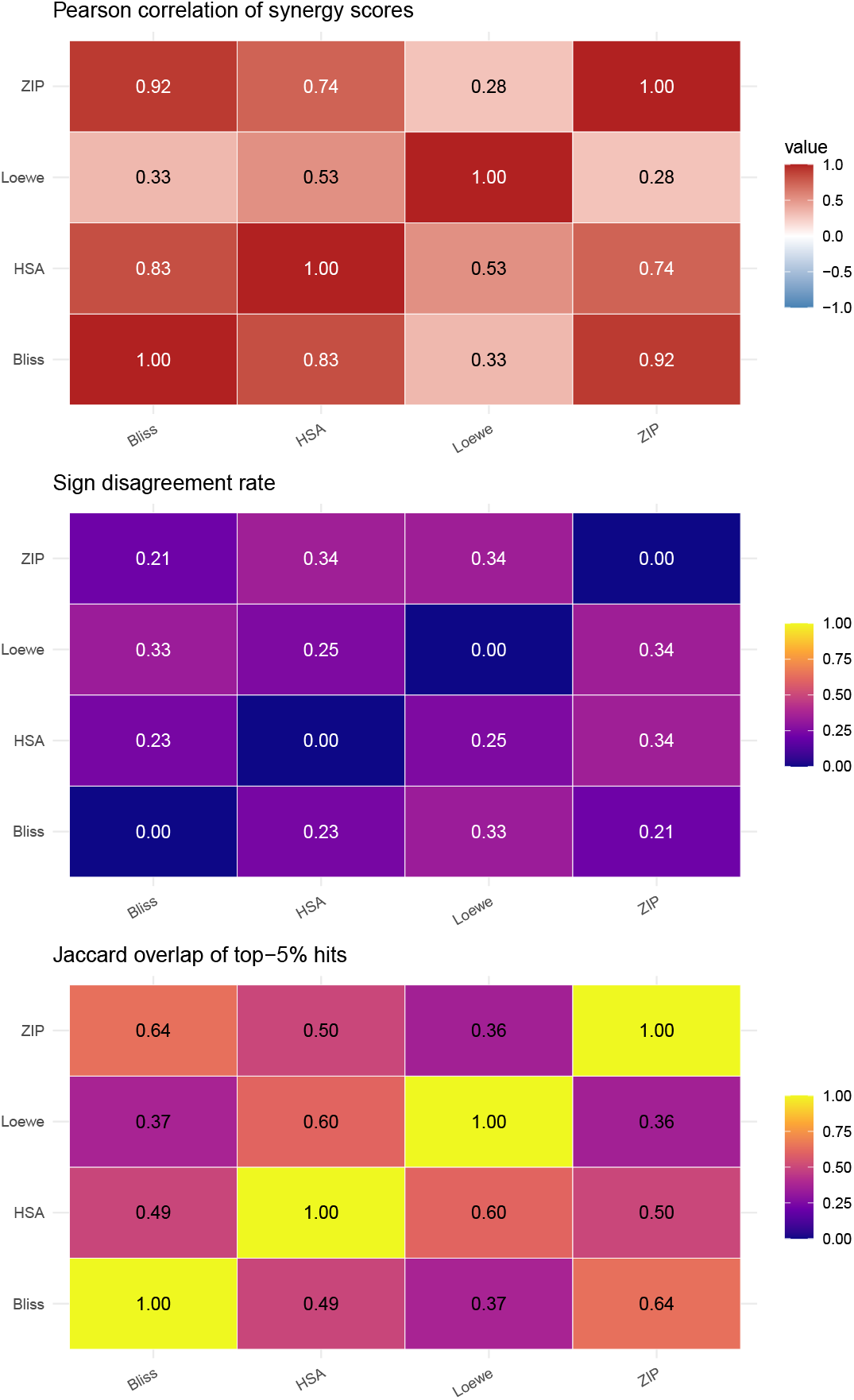
Baseline synergy scores disagree across null models on DrugCombDB. (*n* = 391,652 dose–response matrices with valid scores for all four methods). Top: Pearson correlations of matrix-level summary scores. Middle: fraction of matrices where only one method reports positive synergy. Bottom: Jaccard overlap of top 5% synergy calls. Bliss and ZIP correlate highly (*r* = 0.92), reflecting their shared independence assumptions, whereas Loewe often disagrees with both (*r ≈* 0.3), consistent with its fundamentally different dose-equivalence construction. Disagreement extends to hit calling: the Loewe–ZIP Jaccard overlap among top-5% hits is only 0.36, meaning that which drug pairs are prioritized for follow-up depends heavily on the choice of null model. These results motivate a framework that avoids committing to any single null-model definition of additivity.

### 2.2 A shape-constrained definition of interaction

We model a drug combination experiment as an *I* × *J* grid of responses *Y*_*ij*_ ∈ [0, 1] (viability) measured at increasing doses of two drugs. We transform responses to an unconstrained scale, *Z*_*ij*_ = logit(*Y*_*ij*_), and compute inverse-variance weights from within-cell replicates (Online Methods, *Transform and weights*). The logit transform maps bounded viability data to R, stabilizes variance near the boundaries, and defines additivity on a scale where equal increments correspond to equal log-odds changes in cell survival. We then fit two nested model classes, both solved as convex quadratic programs (Online Methods, *Model classes and estimation*): (i) a flexible monotone surface 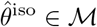 via 2D isotonic regression (the fitted value at each dose pair is constrained to be non-increasing in each drug’s dose); and (ii) a monotone-additive null surface 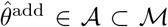, *θ*_*ij*_ = *α* + *u*_*i*_ + *v*_*j*_, where *u*_*i*_ and *v*_*j*_ are each constrained to be monotone non-increasing, so that the combined effect is additive with each drug contributing independently. Their difference defines the *interaction surface*,

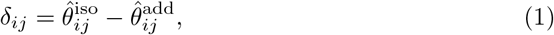

interpretable as synergy (for viability) when *δ*_*ij*_ *<* 0 (lower viability than additive expectation). Importantly, the interaction surface is not constrained to be monotone; it can take any sign pattern across the grid, capturing spatially heterogeneous interaction (e.g., synergy at some dose pairs and antagonism at others). We summarize the overall strength of interaction across the grid by the *interaction energy* 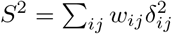 a single scalar that measures weighted squared departure from additivity, analogous to a goodness-of-fit statistic comparing the additive model to the unconstrained monotone model (Online Methods, *Interaction surface and summaries*). We choose *S*^2^ over alternatives such as the difference in residual sums of squares (SSE_add_ − SSE_iso_) because *S*^2^ depends only on the interaction surface *δ* rather than explicitly on residual magnitudes, making it more comparable across datasets in expectation. Note that the interaction signal naturally concentrates at intermediate dose combinations, where one drug is effective enough to reveal synergy or antagonism with the other; at the lowest and highest dose corners, both models predict similar values and *δ* is typically near zero. A key advantage of this definition is that the null model 𝒜 is a nested submodel of the alternative ℳ, both within the same monotone function class, so the interaction surface reflects genuine departures from additivity rather than artifacts of comparing models with different structural assumptions. Figure 2 illustrates the SIR workflow on a DrugCombDB example, from observed viability through the fitted surfaces to the interaction map and bootstrap p-value.

**Fig. 2.**
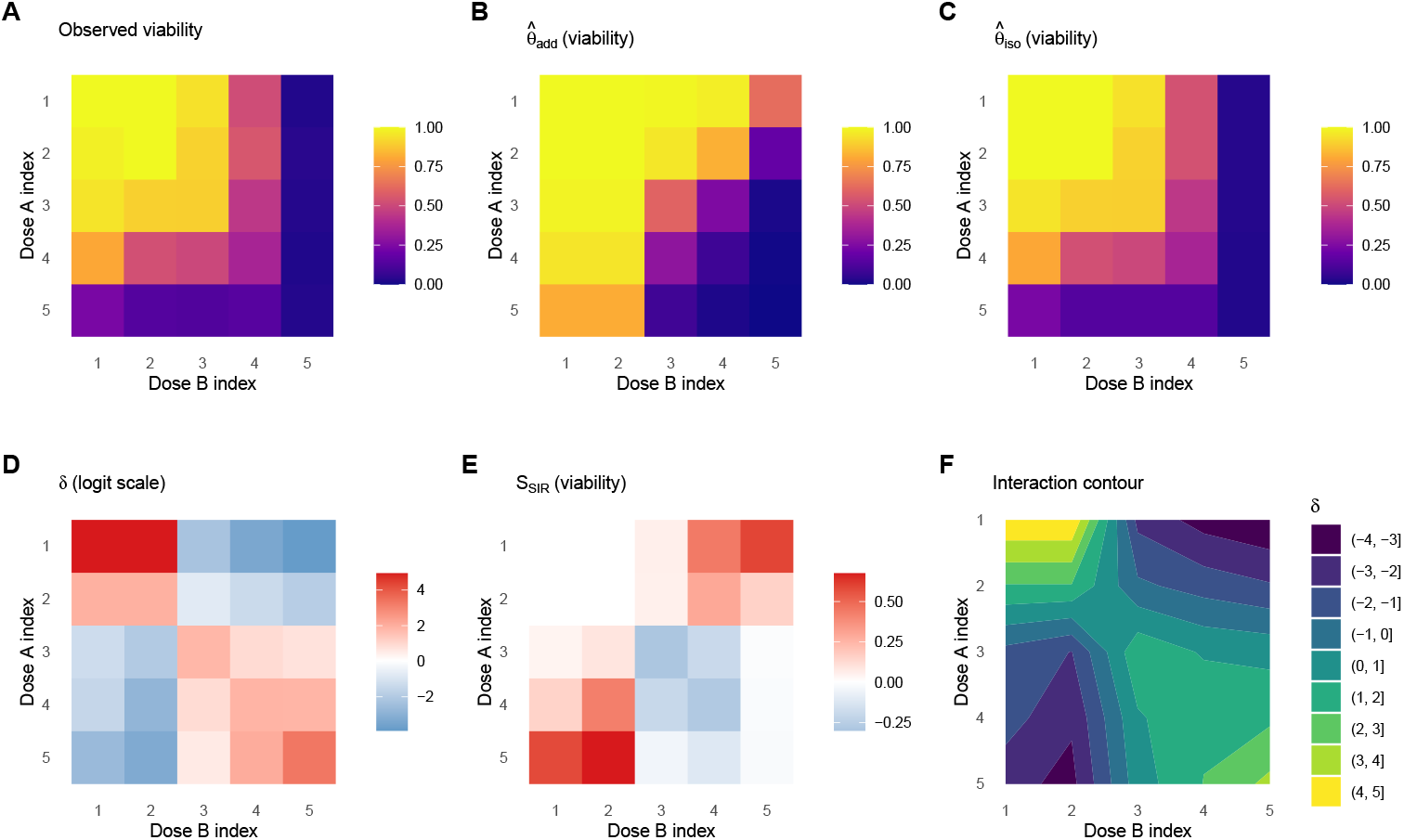
SIR workflow illustrated on a DrugCombDB example. (Gemcitabine + Dinaciclib in EFM192B cells; 5 *×* 5 dose grid). **(A)** Observed viability. **(B)** Monotone-additive null surface 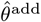 (back-transformed to viability): the best-fitting surface under the assumption of no interaction. **(C)** Unconstrained monotone surface 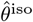 (back-transformed): the best monotone fit without the additivity constraint. **(D)** Interaction surface 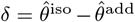 on the logit scale; negative values (blue) indicate synergy, i.e., the combination kills more than the additive model predicts. **(E)** Effect sizes on the viability scale, 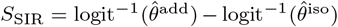. **(F)** Contour map of the interaction surface, showing the spatial pattern of synergy and antagonism across the dose grid. The global interaction test yields *p* = 0.025 (*B* = 200 bootstrap resamples), indicating statistically significant departure from additivity.

### 2.3 Calibrated hypothesis testing by df-corrected wild bootstrap

Existing synergy methods (Bliss, HSA, Loewe, ZIP) return matrix-level synergy scores without p-values, so there is no principled way to assess whether an observed score reflects true interaction or noise, nor to control false-positive rates when screening thousands of drug pairs. SIR addresses this by computing a p-value for each matrix using a wild bootstrap [20, 22] under the monotone-additive null. A standard challenge in bootstrap testing is that residuals from a fitted model underestimate the true noise: because the model was estimated from the same data, it passes closer to the observations than the true underlying dose-response surface would, shrinking the residuals. Bootstrapping with these shrunken residuals produces synthetic datasets that are less variable than real data, biasing the test toward false positives. We correct for this by inflating residuals by a degrees-of-freedom factor before resampling, analogous to the *n/*(*n* − *p*) correction in linear regression variance estimation, where *n* is the number of observed grid cells and *p* is the effective degrees of freedom consumed by the null fit (for a typical 5 × 5 grid with monotone marginals, *n* = 25 and *df*_null_ ≈ 9, giving a ratio of approximately 1.56; Online Methods, *Wild bootstrap inference with df correction*).

To verify that these p-values are well calibrated, we performed a pseudo-null experiment on 300 randomly selected DrugCombDB matrices: for each, we fit the additive null, generated synthetic data by randomly flipping the sign of the corrected residuals (so that no true interaction is present by construction), and reran the full bootstrap test. If the test is properly calibrated, the resulting p-values should follow a uniform distribution, and the false-positive rate at any threshold *α* should be close to *α*. Indeed, the empirical p-values closely follow the uniform distribution (Fig. 3), with observed false-positive rates of 3.3% at *α* = 0.05 and 5.7% at *α* = 0.10. At *α* = 0.05, rejecting 3.3% of true nulls (rather than the nominal 5%) means the test is slightly conservative, a desirable property in screening applications where controlling false positives is more important than maximizing detections. We additionally assessed the test’s statistical power in a controlled simulation study. We generated 8×8 dose-response grids under a monotone-additive null, then injected interaction of increasing strength by adding a localized bump to the surface (Online Methods, *Benchmarks*; Supplementary Methods, *Simulation study design*). Power increases monotonically with interaction strength, rising from near zero at the null to *>*95% at the strongest departures (Fig. 4). These results confirm that SIR has adequate power to detect biologically meaningful interaction while maintaining calibrated false-positive rates.

**Fig. 3.**
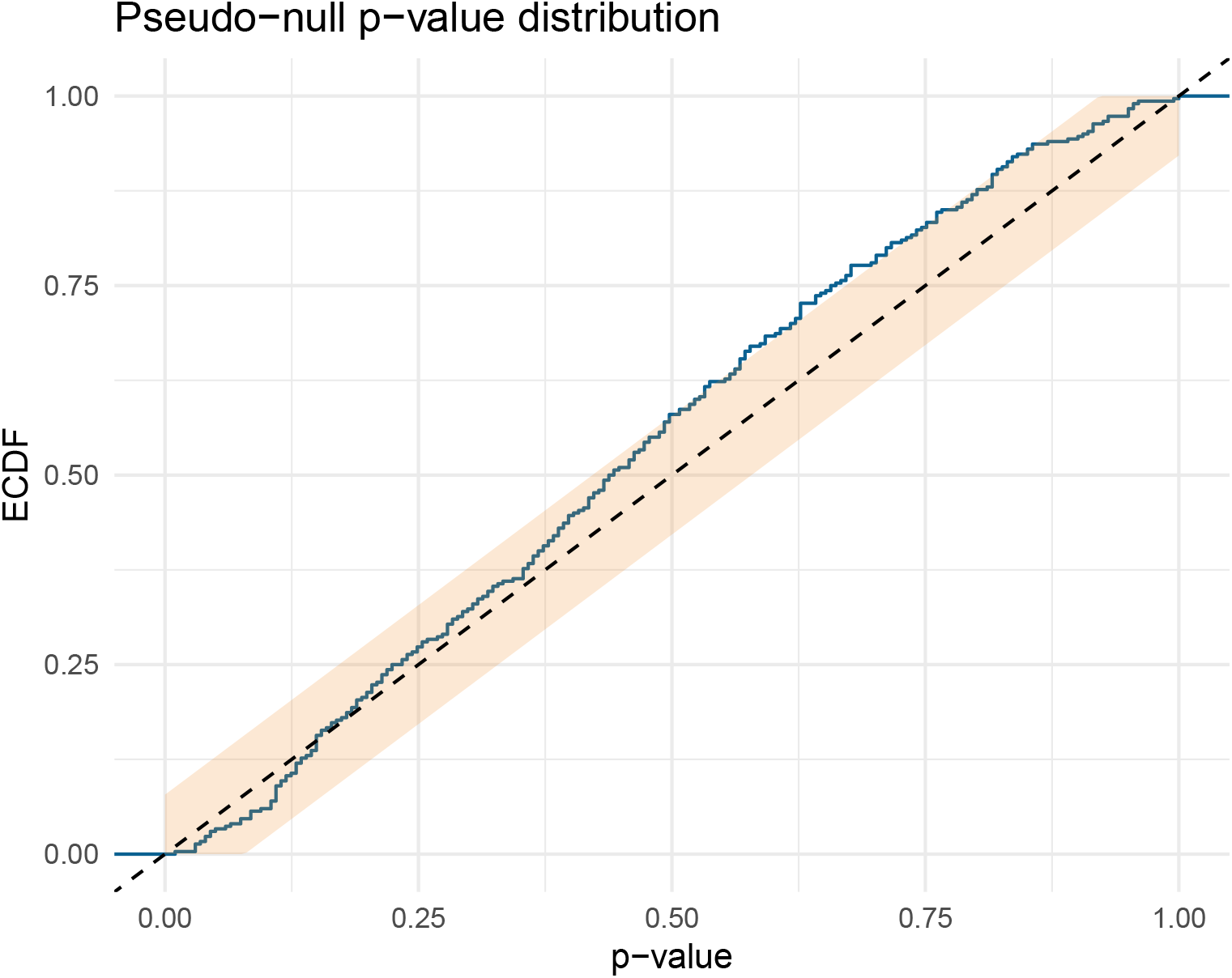
Pseudo-null calibration on DrugCombDB. (*n* = 300 randomly selected matrices; *B* = 200). P-values from the df-corrected wild bootstrap when data are generated under the fitted additive null by sign-flipping corrected residuals, so that no true interaction is present by construction. The dashed diagonal is the Uniform(0, 1) reference expected for a well-calibrated test; the shaded region is a 95% Dvoretzky–Kiefer–Wolfowitz band. The close agreement with the uniform diagonal demonstrates that SIR’s p-values are trustworthy on real screening data with real noise structure, not just in controlled simulations (Fig. 4).

**Fig. 4.**
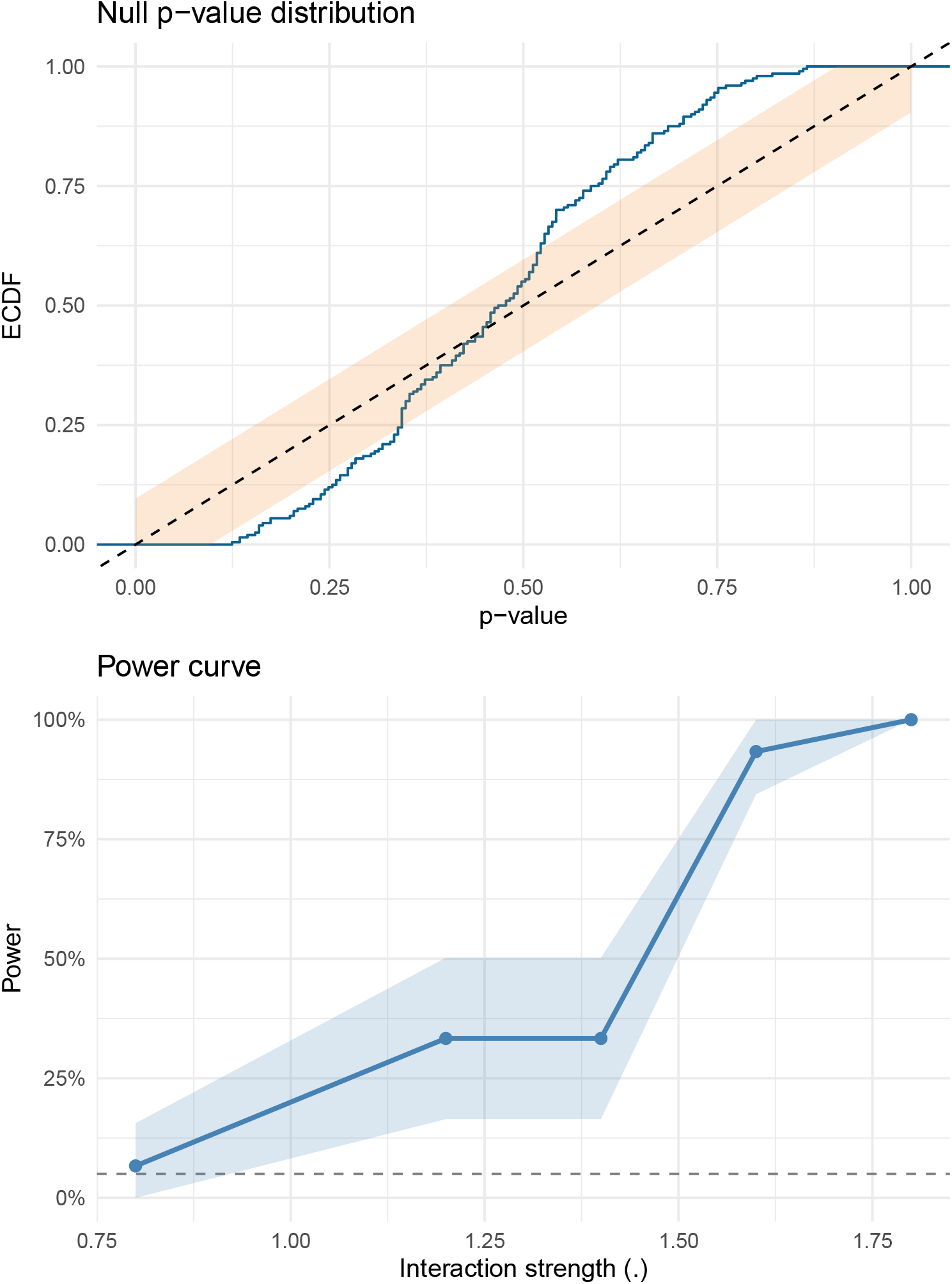
Simulation: calibration and power of the SIR test. (8 *×* 8 grids; *σ* = 0.1 on the logit scale; *B* = 200). Top: empirical cumulative distribution function (ECDF) of p-values under a simulated additive null (*n* = 200 simulations) with 95% Dvoretzky–Kiefer–Wolfowitz band. The CDF closely follows the uniform diagonal, with only minor excursions near the band boundary, confirming that the test does not produce excess false positives. Bottom: power to detect interaction (*α* = 0.05) as interaction strength increases (*n* = 30 simulations per strength). Power rises from near zero at the null to *>*95% at the strongest departures; the steep transition is typical of power curves and reflects the signal crossing the detection threshold.

### 2.4 Higher reproducibility and zero failures on replicate experiments

Reproducible synergy estimates are essential for translating screens into follow-up experiments. If a method assigns different synergy labels to two independent measurements of the same drug pair in the same cell line, its conclusions cannot be trusted. Because these are independent experiments with distinct biological noise, we do not expect identical surfaces, but a reliable method should produce highly correlated interaction patterns. In DrugCombDB, 1,209 unique drug-pair/cell-line combinations were measured in two or more independent experiments, yielding 1,839 replicate pairs (some combinations had three or more experiments, producing multiple pairs). For each replicate pair, we computed the SIR interaction surface *δ* and the corresponding baseline synergy surfaces, then measured the Pearson correlation between replicate surfaces as a measure of reproducibility (Online Methods, *Benchmarks*). SIR’s interaction surface *δ* on the logit scale is substantially more reproducible than all baselines: median replicate correlation is 0.91, compared to 0.53 (Bliss), 0.61 (HSA), 0.74 (Loewe) and 0.71 (ZIP) (Fig. 5). The viability-scale effect sizes *S*_SIR_ are similarly reproducible (median 0.88). The higher reproducibility likely reflects the regularizing effect of the monotonicity constraint: by enforcing a biologically motivated shape, SIR suppresses noise-driven fluctuations that inflate variability in unconstrained pointwise scores.

**Fig. 5.**
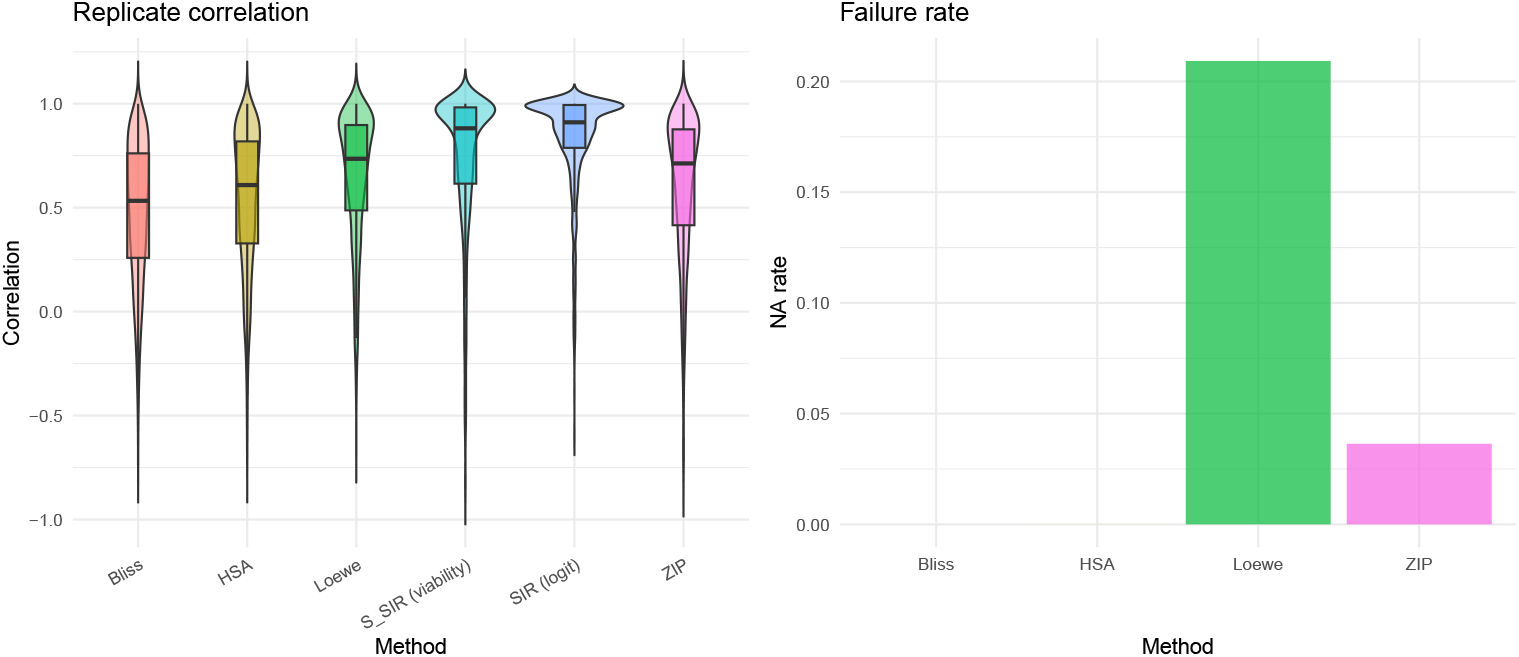
Replicate concordance and failure rates on DrugCombDB. (1,839 replicate pairs from 1,209 experiments). Left: replicate correlation of synergy/interaction surfaces for each method. SIR’s interaction surface on the logit scale achieves a median replicate correlation of 0.91; the viability-scale effect sizes (*S*_SIR_) achieve 0.88. Both are substantially higher than Bliss (0.53), HSA (0.61), ZIP (0.71), or Loewe (0.74), indicating that SIR’s interaction surfaces are more consistent when the same drug pair is measured independently. Right: failure rate (fraction of experiments returning NA/non-finite values). Loewe fails on 20.9% and ZIP on 3.6% of experiments due to marginal curve-fitting failures; SIR never fails because isotonic regression always has a solution.

In addition, parametric baselines can be entirely undefined: Loewe and ZIP fail (non-finite output) in 20.9% and 3.6% of experiments, respectively, while SIR succeeds in all cases (Fig. 5). These failures occur when the marginal dose-response curves do not converge or produce degenerate parameter estimates, a problem that is absent in SIR because isotonic regression is a convex projection that always has a unique solution. For large-scale screens where even a small failure rate can affect thousands of matrices, this robustness is a practical necessity.

### 2.5 A generative surface model enables prediction of missing wells

Drug combination matrices are often incomplete due to plate layout constraints, assay failures, or adaptive designs. Standard synergy scores cannot handle missing entries because they are computed independently at each dose pair; a missing well means a missing score. Because SIR fits an explicit monotone surface to all available data, it naturally predicts unobserved dose pairs by evaluating 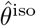 at the missing coordinates and transforming back to viability. In a holdout benchmark on 200 DrugCombDB matrices, we withheld 20% of interior wells and predicted their viabilities from the remaining data. Predictions are accurate (median RMSE 0.040 in viability units), and the induced synergy summaries are similarly stable (median RMSE 0.035 for the cellwise *S*_SIR,*ij*_ values) (Fig. 6). This capability is particularly relevant for adaptive screening designs, where only a subset of dose pairs may be measured in a first pass, and the fitted surface can guide selection of follow-up wells. It also means that SIR can produce synergy summaries even for matrices with irregular or non-rectangular dose layouts, a setting where pointwise baselines would require imputation or exclusion of missing entries.

**Fig. 6.**
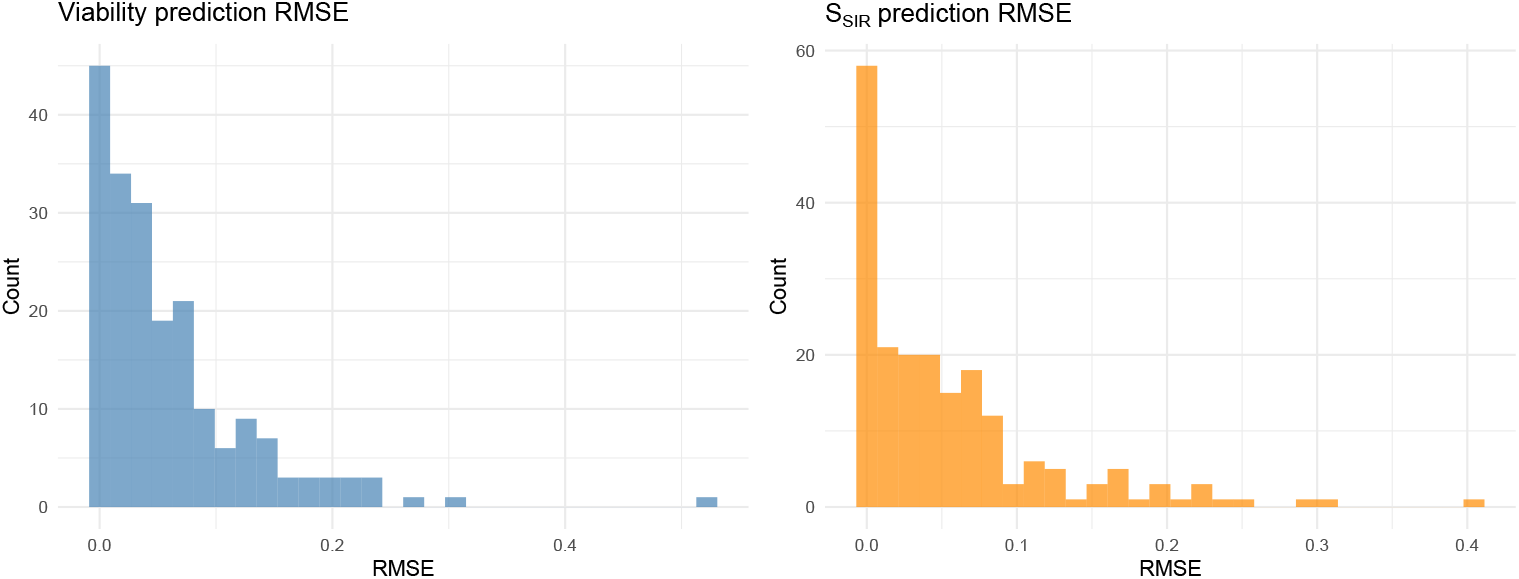
Prediction accuracy under missingness on DrugCombDB. (*n* = 200 matrices, 20% interior wells held out). Left: holdout RMSE for viability predictions. Right: holdout RMSE for the SIR synergy summary *S*_SIR_. Median prediction error is 0.040 in viability units and 0.035 for *S*_SIR_, indicating that the monotone surface captures the underlying dose-response relationship well enough to impute missing measurements. This capability is unique to SIR: existing synergy scores are computed pointwise and cannot predict unobserved dose pairs.

## 3 Discussion

SIR is a statistical framework for drug combination interaction that replaces fragile parametric curve fitting with shape-constrained regression and supplies calibrated p-values by a df-corrected wild bootstrap. The method addresses three recurrent problems in synergy analysis (Table 1). First, baseline null models disagree substantially on the same data (Fig. 1), implying that “synergy” labels are not a stable ground truth [12, 14]. Second, baselines that rely on parametric marginal models (Loewe, ZIP [9]) can fail on real matrices, whereas SIR’s isotonic regression [23] always yields a feasible monotone fit (Fig. 5). Third, the absence of uncertainty quantification in common scoring approaches, including the widely used Combination Index [8, 24], prevents error-rate control and principled aggregation across experiments.

**Table 1.** Capabilities of SIR versus common baselines. ✓ = present; – = absent. Baselines provide pointwise scores without uncertainty; some require parametric curve fitting, which can fail on real data.

| Capability | SIR | Bliss | HSA | Loewe | ZIP |
| --- | --- | --- | --- | --- | --- |
| Statistical test<br>(calibrated p-value) | ✓ | – | – | – | – |
| Predicts missing<br>wells | ✓ | – | – | – | – |
| Shape-constrained<br>regularization | ✓ | – | – | – | – |
| No fit failures<br>on real data | ✓ | ✓ | ✓ | – | – |
| Null assumption | Monotone | Indep. | Max effect | Dose equiv. | Indep. +<br>curve fit |

These issues are relevant for machine-learning-guided combination discovery. ML pipelines that predict synergy scores [17–19] inherit label noise when those scores are inconsistent across models or undefined due to fit failures. SIR may provide more defensible training labels by returning calibrated effect sizes and p-values rather than heuristic scores, though a direct comparison of downstream ML performance remains future work.

### Relationship to other frameworks

MuSyC [3] takes a complementary approach: it fits a parametric multi-parameter surface to disentangle efficacy, potency, and cooperativity synergy. By contrast, SIR avoids parametric assumptions entirely and focuses on calibrated statistical testing of whether any interaction is present, rather than decomposing its mechanistic origin. Response surface approaches [13] and software tools such as Combenefit [11] provide visualization and quantification of synergy using Loewe, Bliss, or HSA, but inherit the limitations of those null models and do not provide calibrated statistical tests. The two frameworks (SIR and MuSyC) address different questions and could be used in sequence: SIR to identify interacting pairs with controlled error rates, followed by parametric modeling (MuSyC or others) to characterize the nature of confirmed interactions.

### Direction of interaction and multiplicity

The global interaction energy *S*^2^ is inherently two-sided: it tests whether the doseresponse surface deviates from monotone additivity anywhere, without committing to a single sign across the grid. After rejecting the global null, users may summarize direction using directional energies (Supplementary Methods, *Global and directional summaries*) and, if desired, test whether the interaction is predominantly synergistic or antagonistic. Because this directional test is only performed after the global test has already confirmed that interaction exists, it does not require a separate multiple-testing correction: the global test acts as a gatekeeper that controls the overall false-positive rate.

### Monotonicity as a minimal assumption

Monotonicity can fail in cases such as hormesis or biphasic responses [15]. Rather than assuming a parametric form, we treat monotonicity as a regularizing constraint, a weak assumption that stabilizes estimation on sparse, noisy grids without imposing a specific functional form. Dataset-scale diagnostics on DrugCombDB indicate that most apparent monotonicity violations are small in magnitude and consistent with noise (Supplementary Fig. 1), supporting monotone regression as a pragmatic default. When strong non-monotonicity is suspected, diagnostics can flag affected matrices for alternative modeling.

### Interaction at dose extremes

The interaction surface *δ* naturally concentrates at intermediate dose combinations: at the lowest doses, neither drug is effective and both models predict viability near one (a ceiling effect); at the highest doses, both drugs independently drive viability toward zero, leaving little room for additional combinatorial effect (a floor effect). This is pharmacologically expected and does not bias the global test, because *S*^2^ sums over all grid cells and near-zero *δ* values at the corners contribute negligibly. The logit transform partially mitigates the compression by stretching both tails, but the floor effect persists in practice. Users interested in dose-specific synergy should examine the interaction surface directly rather than relying solely on the global test.

### Choice of transform

The null hypothesis of monotone additivity is defined on the scale determined by the response transform. Because different transforms (logit, identity, asinh) impose different geometries, inference results can depend on this choice, a property shared by all parametric and semiparametric interaction tests. We recommend the logit link as a principled default for bounded viability data: it maps [0, 1] to ℝ, stabilizes variance near the boundaries, and defines additivity on a scale where equal increments correspond to equal log-odds changes. When the appropriate scale is uncertain, we recommend reporting results under two or more transforms as a sensitivity check (Supplementary Note 2).

### Limitations

SIR tests for departure from monotone additivity but does not decompose the interaction into mechanistic components (e.g., potency versus efficacy synergy, as in MuSyC [3]). The monotone-additive null treats each drug’s contribution as a fixed monotone function, which may be overly flexible for low-dimensional grids (e.g., 3 × 3) where few data points are available for estimation; in such settings, the test may have limited power. The wild bootstrap assumes that errors are independent across grid cells, which could be violated if systematic plate effects or spatial correlations are present. Finally, while the logit transform is a principled default, the choice of transform affects the null hypothesis and therefore the results, and there is no universally correct scale for defining additivity.

### Practical guidance

For exploratory screens with thousands of matrices, we recommend *B* = 200 bootstrap resamples with Benjamini–Hochberg FDR correction on the resulting p-values. For confirmatory studies focused on a small number of candidate combinations, *B* should be increased to 1,000 or more to achieve finer p-value resolution. Grid sizes of 4 × 4 or larger provide sufficient degrees of freedom for the monotone-additive fit; smaller grids may benefit from reduced models. When within-cell replicates are available, inverse-variance weighting improves efficiency by down-weighting noisy dose pairs. SIR’s fitted surfaces and interaction maps can be visualized alongside the scalar p-value to provide spatial insight into where on the dose grid interaction is strongest.

### Outlook

The framework readily extends to alternative response transforms, weighted designs, and different global statistics. SIR is, to our knowledge, the first synergy-testing framework that combines isotonic regression, a monotone-additive null projection, and calibrated bootstrap p-values for drug combination screens. More broadly, replacing heuristic scores with statistically grounded estimators can help reconcile the fragmented landscape of synergy models [13, 15, 25] by enabling calibrated discovery and reproducible benchmarking. As a generalizability check, we replicated the baseline disagreement and pseudo-null calibration analyses on NCI-ALMANAC [26], an independent large-scale combination screen; both findings held (Supplementary Figs. 2–3). Looking forward, SIR could be extended to higher-order combinations (three or more drugs) by defining monotone-additive null models on higher-dimensional dose grids, though computational cost and the curse of dimensionality would need to be addressed. Integration with downstream machine learning pipelines, where SIR’s calibrated p-values and interaction surfaces replace heuristic synergy scores as training labels, is another promising direction.

## 4 Online Methods

### 4.1 Data sources and preprocessing

We evaluated the method primarily on DrugCombDB [21], a curated database of drug combination experiments with dose matrices across many drugs and cell lines. We used viability responses scaled to [0, 1] and standardized doses to a common unit when available. DrugCombDB provides baseline synergy scores for Bliss, HSA, Loewe and ZIP via SynergyFinder conventions [4]; we used these for baseline disagreement analyses.

### 4.2 Transform and weights

Let *Y*_*ij*_ ∈ [0, 1] denote viability at dose pair (*i, j*). We transform responses to *Z*_*ij*_ = logit(*Y*_*ij*_), clamping *Y*_*ij*_ to [*ϵ*, 1 − *ϵ*] with *ϵ* = 10^−6^ to avoid infinities. We use the logit link because viability is bounded in [0, 1] and the logit stabilizes variance near the boundaries; alternative transforms (identity, asinh, log) are explored in Supplementary Note 2. When replicate measurements are available, we compute inverse-variance weights

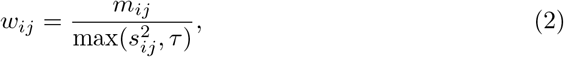

where *m*_*ij*_ is the number of replicates, 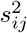 is the sample variance, and *τ* = 10^−6^ is a variance floor that prevents degenerate weights. Weights are Winsorized at the 99^th^ percentile to limit the influence of extremely low-variance cells. When no replicates are available (*m*_*ij*_ = 1), all weights default to 1*/τ* (i.e., uniform weighting).

### 4.3 Model classes and estimation

Let ℳ denote the set of monotone surfaces on an *I* × *J* grid (non-increasing in each coordinate for viability). The isotonic estimator is the weighted least-squares projection

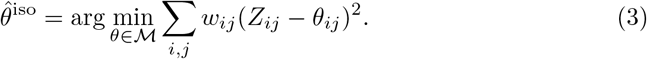

Let 𝒜 ⊂ ℳ denote the monotone-additive class *θ*_*ij*_ = *α* + *u*_*i*_ + *v*_*j*_, with *u* and *v* constrained to be monotone in the same direction and with identifiability constraints (*u*_1_ = 0, *v*_1_ = 0) to fix the intercept and prevent trading constants between *α, u*, and *v*. We compute 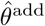 as the weighted least-squares projection onto 𝒜. Both problems are solved as convex quadratic programs with linear inequality constraints using OSQP [27].

### 4.4 Interaction surface and summaries

The interaction surface 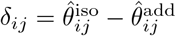 is defined on the logit scale. To report effect sizes in interpretable viability units, we back-transform each surface and take their difference:

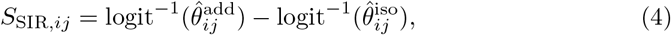

so that positive values of *S*_SIR_ indicate synergy (the combination kills more than the additive prediction). Global interaction strength is summarized by the interaction energy 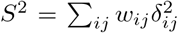. For the bootstrap test, we use the normalized variant *T* = *S*^2^*/* ∑_*ij*_ *w*_*ij*_, which adjusts for grid size and weighting.

### 4.5 Wild bootstrap inference with df correction

To test the null hypothesis of monotone additivity, we use a Rademacher wild bootstrap [20]. We fit the null model 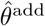 and compute residuals 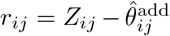. Because the fitted null absorbs some noise, residuals underestimate the true error variance. We correct for this by inflating residuals:

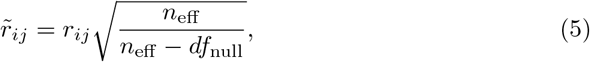

where *n*_eff_ is the number of grid cells with finite, positively weighted observations, and *df*_null_ is the effective degrees of freedom consumed by the monotone-additive fit. Unlike linear regression, where df equals the fixed number of parameters, isotonic regression pools adjacent dose levels that violate monotonicity into tied groups, so its effective df depends on the data: if the observed marginal responses are already monotone, each dose level retains its own fitted value and df is large; if they are highly non-monotone, many levels are pooled and df is small. We approximate *df*_null_ by counting the number of distinct fitted levels in the monotone main effects *û* and 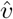 (see Supplementary Methods, *Degrees-of-freedom correction in the wild bootstrap*). We then generate *B* bootstrap datasets 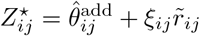, where *ξ*_*ij*_ ∈ {−1, +1} are independent Rademacher random variables [20, 22], refit both models on each *Z*^⋆^, and recompute the test statistic 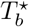. The p-value is

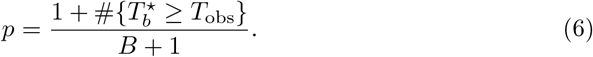

The default *B* = 200 provides adequate resolution for exploratory screens (p-value increments of ≈0.005); for confirmatory analyses requiring stringent thresholds (e.g., *α <* 0.01 after multiple-testing correction), *B* should be increased to 1,000–5,000. On a single core, the full test takes ≈0.9 s per 4 × 4 matrix and ≈1.5 s per 10 × 10 matrix with *B* = 200; a 400,000-matrix screen completes in ≈14 hours on 8 cores.

### 4.6 Multiple testing in screens

When applying the test across a screen of *M* matrices, standard false-discovery-rate procedures (e.g., Benjamini–Hochberg [28]) can be applied directly to the per-matrix p-values. Users should note that the smallest achievable p-value is 1*/*(*B* + 1); for large screens where the FDR threshold may require very small p-values, increasing *B* is recommended. Stratifying by cell line or plate before applying FDR correction can improve power, because matrices within the same cell line or plate tend to share similar noise levels, making the p-value distribution within each stratum more homogeneous.

### 4.7 Benchmarks

**Baseline disagreement:** We computed pairwise correlations, sign disagreement rates, and top-hit overlaps across baseline scores on DrugCombDB matrices with valid (non-NA) Bliss, HSA, Loewe and ZIP scores (391,652 matrices). **Replicate concordance:** We sampled repeated experiments for the same drug pair and cell line, computed synergy/interaction surfaces for each replicate experiment, and evaluated replicate correlations and failure rates. **Missingness prediction:** For each of 200 matrices, we held out 20% of interior wells, fit the SIR model on the remaining wells, and evaluated RMSE on held-out viabilities and derived synergy summaries. **Pseudo-null calibration:** We generated pseudo-null data by sign-flipping df-corrected residuals from the fitted null and recomputed bootstrap p-values. **Simulation:** We constructed monotone-additive null surfaces on 8×8 grids (intercept plus monotone row and column effects), then injected interaction by adding a localized bump projected onto the monotone cone to ensure model-consistent alternatives (see Supplementary Methods, *Simulation study design*). Gaussian noise (*σ* = 0.1 on the logit scale) was added to generate observed data. Under the null (no injected interaction), we evaluated Type I error calibration by running *n* = 200 simulations and checking that p-values are uniformly distributed (Fig. 4, top). Under the alternative (injected interaction at five positive strengths), we evaluated power by running *n* = 30 simulations per strength and measuring rejection rate at *α* = 0.05 (Fig. 4, bottom). All simulations used *B* = 200 bootstrap resamples.

## 4.8 Code availability

The SIR R package and all analysis scripts are available at https://github.com/AsiaeeLab/SIR (Supplementary Note 1). The repository includes source code, configuration files, experiment notebooks, and instructions to reproduce all figures and tables in this manuscript.

## 4.9 Data availability

DrugCombDB data are publicly available at http://drugcombdb.denglab.org/ [21]. NCI-ALMANAC data are available from the National Cancer Institute at https://wiki.nci.nih.gov/display/NCIDTPdata/NCI-ALMANAC. Download instructions, checksums, and file placement details are provided in the repository README.

## Acknowledgements

A.A. and S.P. were supported in part by the Patient-Centered Outcomes Research Institute (PCORI) award ME-2023C1-32148. A.A. was additionally supported by the National Human Genome Research Institute (NHGRI) of the National Institutes of Health under award R00 HG011367. J.P.L. received support from the National Cancer Institute and the National Center for Advancing Translational Sciences of the NIH (P50CA127001-16, CCSG P30CA016672-46, and CCTS UM1TR004906). The content is solely the responsibility of the authors and does not necessarily represent the official views of PCORI, the NIH, or any of the funding agencies listed above.

## Author contributions

A.A. and K.R.C. conceived the method. A.A. developed the statistical framework, implemented the software, and wrote the manuscript. J.P.L. contributed to the statistical methodology and interpretation. S.P. edited the manuscript. H.H.P. provided biological context and interpretation. K.R.C. supervised the project. All authors reviewed and approved the final manuscript.

## Supplementary Information

### Supplementary Methods

#### 1 Model-consistent definition of interaction

This section provides the formal definitions underlying the SIR interaction surface introduced in the main text. The key idea is that both the null model (no interaction) and the alternative model (unrestricted interaction) are defined within the same class of monotone functions, so that any detected interaction reflects a genuine departure from additivity rather than an artifact of comparing models with different structural assumptions.

Let *Z* ∈ ℝ^*I×J*^ denote transformed responses on an *I* × *J* dose grid and *w*_*ij*_ ≥ 0 denote weights. Let ℳ be the set of monotone surfaces (for viability, non-increasing in each coordinate) and let 𝒜 ⊂ ℳ be the monotone-additive class *θ*_*ij*_ = *α*+*u*_*i*_+*v*_*j*_, where *u*_*i*_ and *v*_*j*_ are each constrained to be monotone non-increasing, with identifiability constraints *u*_1_ = 0 and *v*_1_ = 0 to prevent trading constants between the intercept *α* and the marginal effects. We define the isotonic and additive estimators as weighted projections:

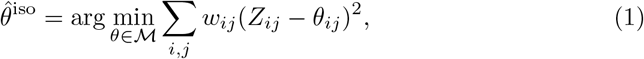

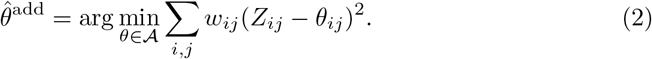

The interaction surface is

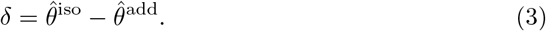

This definition is *model-consistent* : interaction is defined as the component of the best monotone fit that cannot be explained by the best monotone-additive fit, within the same constrained geometry.

##### Why the weighted mean interaction is zero

Both 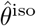 and 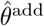 are translation-invariant projections: if *θ* ∈ ℳ (or 𝒜), then *θ* + *c***1** is feasible for any constant *c*. With a squared-error objective, translation invariance implies that both projections preserve the weighted mean of *Z*, and therefore ∑_*ij*_ *w*_*ij*_*δ*_*ij*_ = 0. Consequently, interaction can be mixed-sign even for strongly synergistic combinations; directional summaries should therefore avoid relying on the signed mean of *δ*.

#### 2 Global and directional summaries

##### Global interaction energy

We summarize interaction magnitude with the weighted energy

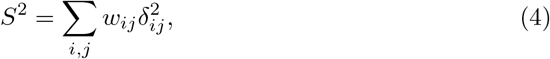

and report either the normalized mean energy *S*^2^/∑*w* or a studentized version *S*^2^*/*SSE_add_ for inference, where 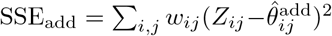. The default test statistic throughout this paper is *T* = *S*^2^*/* ∑_*ij*_ *w*_*ij*_; the studentized alternative is noted for completeness but is not used in any reported results.

##### Directional energies and a bounded direction index

For viability, synergy corresponds to *δ*_*ij*_ *<* 0. Define one-sided energies

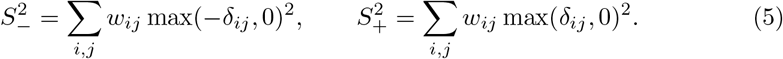

We summarize direction by the bounded index

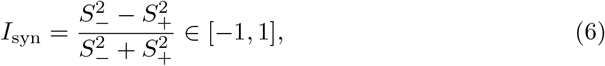

where values near +1 indicate predominantly synergy-like interaction (negative deviations for viability), values near −1 indicate antagonism-like interaction, and values near 0 indicate mixed-sign interaction.

##### Sequential inference and multiplicity

The global test based on *S*^2^ (or *S*^2^*/*SSE_add_) answers “does interaction exist?” and is inherently two-sided. Directional inference can be performed after rejecting the global null using a step-down procedure: first test interaction at level *α*; only if rejected, test direction using a pre-specified directional statistic (e.g., *I*_syn_ or 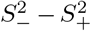) with an appropriate bootstrap reference. Because directional testing is only performed after the global test has confirmed that interaction exists, the global test acts as a gatekeeper: it controls the overall false-positive rate without requiring a separate multiple-testing correction for the directional step. This is analogous to only examining individual regression coefficients after a global F-test has rejected the null that all coefficients are zero.

#### 3 Degrees-of-freedom correction in the wild bootstrap

The wild bootstrap generates pseudo-data under the fitted null:

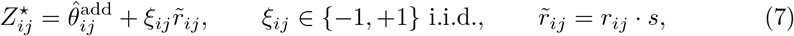

where 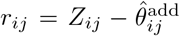 are null residuals and *s* is a residual scaling factor. Because 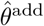 is estimated from the same data, residuals are shrunk relative to the true noise variance. We therefore inflate residuals by a degrees-of-freedom (df) factor

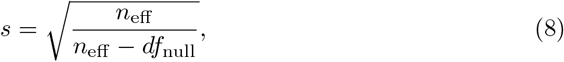

where *n*_eff_ is the number of finite, positively weighted grid cells and *df*_null_ is the effective degrees of freedom consumed by the monotone-additive fit.

Unlike linear regression, where degrees of freedom equals the fixed number of parameters, isotonic regression pools adjacent dose levels that violate monotonicity into tied groups, so its effective df depends on the data. If the observed marginal responses are already monotone, each dose level retains its own fitted value and df is large; if they are highly non-monotone, many levels are pooled and df is small. We approximate *df*_null_ by counting the number of distinct fitted levels in the monotone main effects *û* and 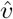. This approximation is analogous to counting the number of free parameters in a piecewise-constant fit: each group of dose levels that share the same fitted value acts as one free parameter, so the total df is approximately the number of such groups across both margins. See [1] for wild bootstrap foundations.

#### 4 Simulation study design

We constructed simulated dose-response data as follows.

##### Null surface generation

For each simulation replicate, we drew monotone row effects *u*_1_ ≥ *u*_2_ ≥ · · · ≥ *u*_8_ and column effects *v*_1_ ≥ *v*_2_ ≥···≥ *v*_8_ independently from standard normal distributions and then sorted them to enforce monotonicity. An intercept *α* was drawn from Uniform(1, 3) to place the surface at biologically plausible viability levels. The null surface is 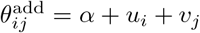 on the logit scale.

##### Interaction injection

To generate alternatives, we added a localized bump centered at the middle of the 8 × 8 grid. The bump amplitude at each cell is *A* exp (−∥ (*i, j*) − center∥^2^*/*(2*σ*^2^), where *A* is the interaction strength parameter, center = (4.5, 4.5), and *σ* = 2. The bump is then projected onto the monotone cone to ensure that the resulting surface remains feasible under the alternative model; this makes the alternative *model-consistent* (interaction is detectable by 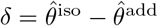).

##### Observed data

Gaussian noise 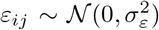 with *σ*_*ε*_ = 0.1 (on the logit scale) was added to each cell of the surface to generate observed *Z*_*ij*_. No replicates were simulated (*m*_*ij*_ = 1, uniform weights).

##### Simulation grid

Interaction strengths *A* ∈ {0, 0.8, 1.2, 1.4, 1.6, 1.8} were used. At *A* = 0 (the null), we ran *n* = 200 independent replicate simulations to obtain a precise empirical CDF for the Type I error check (Fig. 4, top panel, in the main text). At each of the five positive strengths (*A >* 0), we ran *n* = 30 replicates and measured the rejection rate at *α* = 0.05 to estimate power (Fig. 4, bottom panel). All simulations used *B* = 200 bootstrap resamples.

#### 5 Dataset-scale monotonicity diagnostics

SIR assumes that the dose-response surface is monotone (non-increasing in each drug’s dose for viability endpoints). This assumption is biologically motivated by the expectation that higher drug concentrations should not decrease killing, but it may be violated in practice by measurement noise or genuine non-monotone phenomena such as hormesis. To assess how often and how severely monotonicity is violated in real screening data, we computed diagnostic statistics on DrugCombDB.

##### Diagnostic definition

For each matrix, we check every pair of adjacent dose levels along each drug’s axis (holding the other drug’s dose fixed) and record whether viability increases rather than decreases with dose. For a 5 × 5 grid, there are (5 − 1) × 5 = 20 such adjacent comparisons per axis; for a 4 × 4 grid, (4 − 1) × 4 = 12. We report the fraction of these comparisons that violate monotonicity along drug A (*f*_*A*_) and drug B (*f*_*B*_), as well as the maximum magnitude of any violation (*m*) in viability units. We summarize each matrix by *f*_joint_ = (*f*_*A*_ + *f*_*B*_)*/*2 and *m* = max(*m*_*A*_, *m*_*B*_).

**Supplementary Fig. 1.**
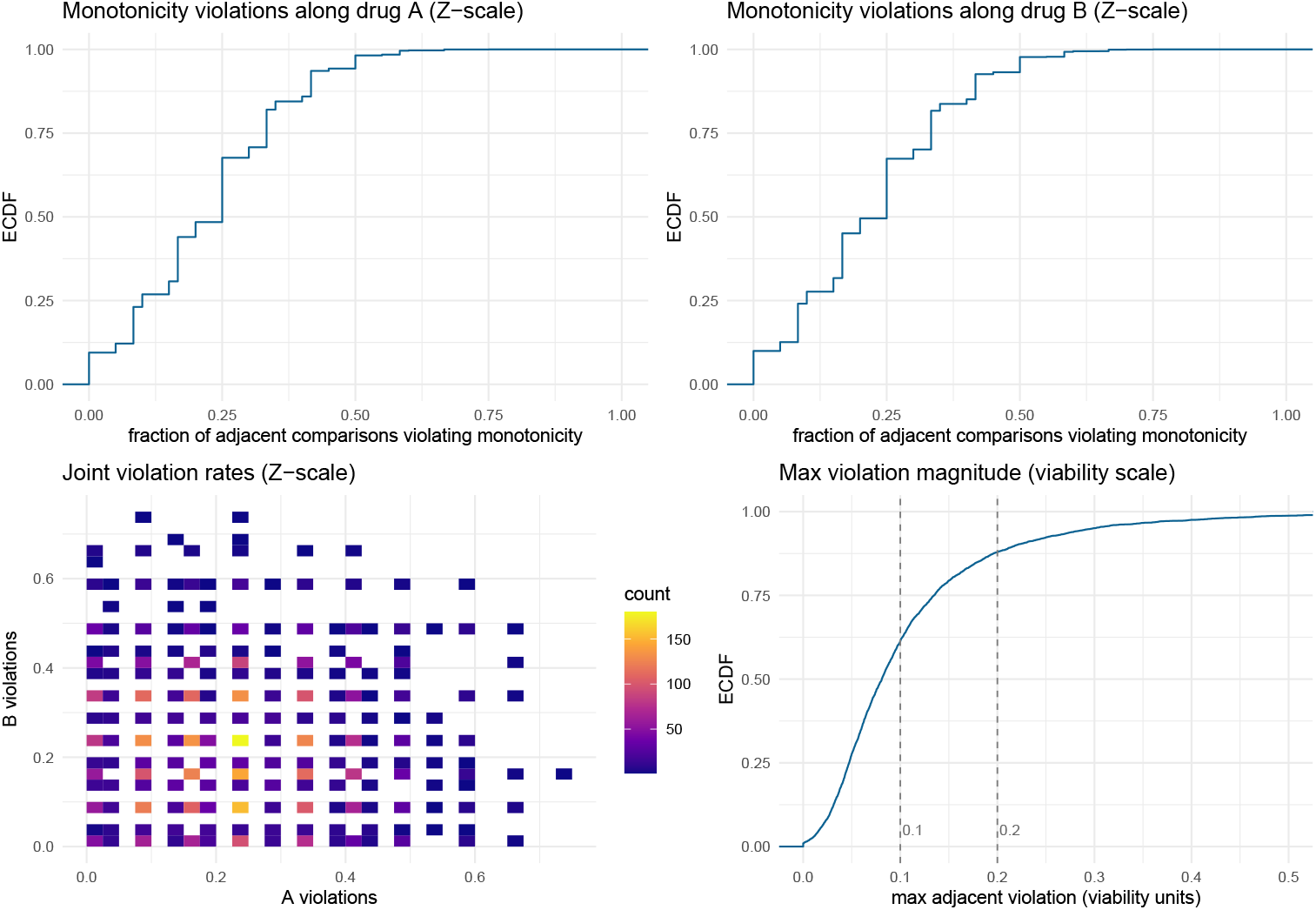
Monotonicity violation frequency and magnitude on Drug-CombDB. (5,000 randomly sampled matrices). Top row: empirical CDFs of the fraction of adjacent dose comparisons that violate monotonicity along each drug’s dose axis. Bottom left: joint distribution of violation fractions across both axes. Bottom right: empirical CDF of the maximum adjacent violation magnitude on the viability scale; dashed lines at 0.1 and 0.2 viability units for reference. Violations are frequent (median fraction *≈*0.25 per axis) but small in magnitude (median maximum violation 0.081; 90th percentile 0.219), consistent with measurement noise rather than systematic non-monotone biology. These results support monotone regression as a pragmatic default; the small fraction of matrices with large violations (*<*5% exceed 0.3) can be flagged for inspection or alternative modeling.

##### Empirical results on DrugCombDB

On a random sample of 5,000 DrugCombDB matrices (3,538 of size 4 × 4 and 1,462 of size 5 × 5), the median violation fraction was *f*_*A*_ = *f*_*B*_ = 0.25, meaning roughly one in four adjacent comparisons shows a reversal. In concrete terms, a typical 5 × 5 matrix has about 5 violations out of 20 adjacent pairs per axis, and a typical 4 × 4 matrix about 3 out of 12. Crucially, these violations are small in magnitude: the median maximum adjacent violation was only 0.081 in viability units (90th percentile 0.219; 95th percentile 0.298), and 61.2% (87.9%) of matrices had maximum adjacent violation ≤ 0.1 (≤ 0.2). This pattern is consistent with measurement noise rather than systematic non-monotone biology: genuine hormesis or biphasic effects would produce large, consistent reversals across multiple dose pairs, not the scattered small fluctuations observed here.

**Supplementary Fig. 2.**
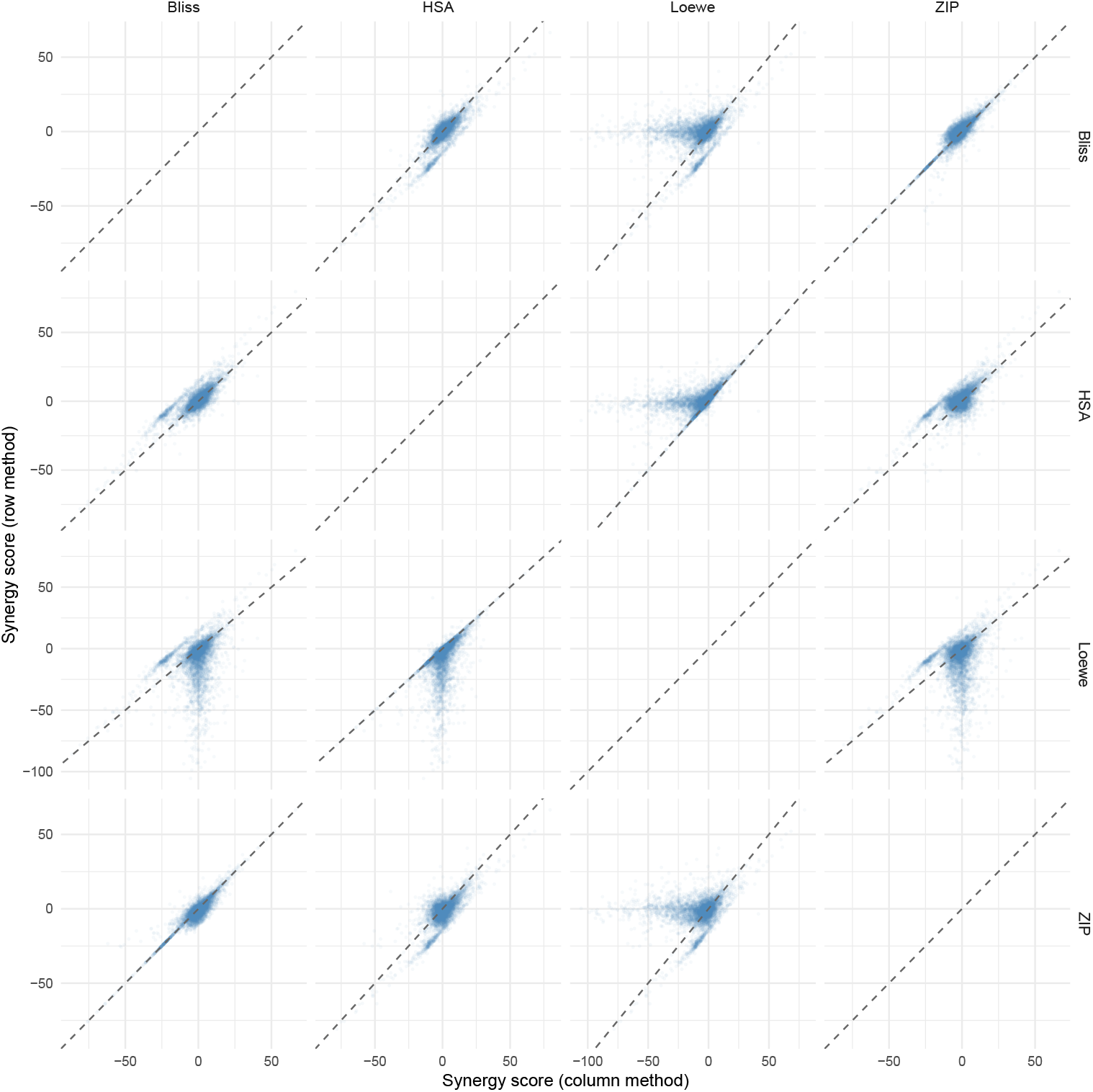
Pairwise scatterplots of baseline synergy scores on DrugCombDB. (5,000 randomly subsampled matrices). Each panel shows one method’s matrix-level summary score against another; the dashed diagonal is the identity line. The near-linear Bliss–ZIP relationship (top right) confirms their high correlation (*r* = 0.92), while the Loewe–ZIP and Loewe–Bliss panels reveal broad clouds with many sign-discordant points (opposite quadrants), explaining the low correlations (*r ≈* 0.3) and high sign disagreement rates reported in Fig. 1 of the main text.

#### 6 Supplementary Discussion: baseline disagreement metrics

To quantify the extent to which different synergy scoring methods agree on which drug combinations are synergistic, we computed three complementary metrics across all DrugCombDB dose-response matrices with valid (non-NA) scores for all four baseline methods.

##### Pearson correlation

For each pair of methods, we computed the Pearson correlation between their matrix-level summary scores across all matrices. High correlation indicates that the two methods rank drug pairs similarly; low correlation indicates that the methods capture fundamentally different aspects of the dose-response landscape or define additivity differently.

##### Sign disagreement rate

For each pair of methods, we computed the fraction of matrices for which exactly one method reports positive synergy (score *>* 0) while the other reports antagonism (score *<* 0). This captures qualitative disagreement: the methods not only rank differently but reach opposite conclusions about the direction of interaction.

##### Top-hit overlap

For each pair of methods, we identified the top *q* = 5% most synergistic matrices by each method’s score, then computed the Jaccard index (intersection over union) between these hit sets. Low overlap means that the two methods would prioritize different drug pairs for experimental follow-up, with direct consequences for resource allocation in screening campaigns.

These summaries complement per-dose-point disagreement by quantifying instability at the level of global hit calling, which is the primary output of most synergy screening workflows.

#### 7 Supplementary Note 1: software and reproducibility

The SIR software and all analysis code are available at https://github.com/AsiaeeLab/SIR. The repository contains the R source code implementing the SIR framework (isotonic regression, monotone-additive fitting, wild bootstrap testing), baseline method implementations (Bliss, HSA, Loewe, ZIP), visualization functions, and the complete set of scripts used to generate every figure and table in this manuscript.

All analyses are driven by YAML configuration files (e.g., `configs/default.yaml`) that specify transform parameters, bootstrap settings, and output directories, ensuring that results are fully reproducible from a single configuration. The constrained regression problems (isotonic and monotone-additive fits) are solved as convex quadratic programs using OSQP [2]. Intermediate results are stored as Parquet files, enabling exact reproduction of reported summary statistics without rerunning the full pipeline. The repository README provides step-by-step instructions for downloading the required datasets, restoring the R environment, and reproducing all results.

#### 8 Supplementary Note 2: transform sensitivity

The null hypothesis of monotone additivity is defined on the scale determined by the response transform. Different transforms (identity on raw viability, logit, asinh, log) induce different notions of “additivity” and therefore different interaction surfaces and p-values. This is not a weakness of SIR but a fundamental property of any interaction test: the notion of “no interaction” depends on the scale.

For example, under the identity transform (raw viability), “additive” means that drug effects sum in viability units: if drug A reduces viability by 0.2 and drug B by 0.3, the combination is expected to reduce it by 0.5. Under the logit transform, “additive” means effects sum on the log-odds scale, which compresses near the boundaries (0 and 1) and produces a different expected combination surface. A drug pair that significantly departs from additivity on one scale may be consistent with additivity on another. On a single example matrix, changing the transform from identity to logit changed the p-value from *p* ≈ 0.01 to *p* ≈ 0.20, not because the method is unstable, but because the two transforms define different null hypotheses and the data are consistent with additivity on the log-odds scale but not on the raw viability scale.

We recommend the logit link as a principled default for viability data because it maps [0, 1] to ℝ, stabilizes variance near the boundaries, and defines additivity on a scale where equal increments correspond to equal log-odds changes in cell survival. When the appropriate scale is uncertain, we recommend reporting results under two or more transforms as a sensitivity analysis. The accompanying code repository provides a detailed exploration of transform effects on fitted surfaces, interaction maps, and bootstrap p-values.

#### 9 NCI-ALMANAC replication

To assess generalizability beyond DrugCombDB, we repeated the baseline disagreement and pseudo-null calibration analyses on NCI-ALMANAC [3], a large-scale combination screen from the National Cancer Institute.

##### Baseline disagreement

On a sample of 1,918 NCI-ALMANAC matrices with valid (non-NA) scores for all four baselines, Bliss and ZIP correlate near-perfectly (*r >* 0.99), while Loewe disagrees strongly with all other methods: sign disagreement rates of 38–43% and top-5% Jaccard overlaps as low as 0.06 (Supplementary Fig. 2). These patterns replicate and amplify the fragmentation observed on DrugCombDB (Fig. 1 of the main text). Notably, Loewe fails (non-finite output) on 97% of NCI-ALMANAC experiments, compared to 20.9% on DrugCombDB, reflecting the more challenging dose–response landscape of this dataset.

##### Pseudo-null calibration

On 120 NCI-ALMANAC matrices subjected to the pseudo-null procedure (sign-flipping df-corrected residuals, *B* = 200), the resulting p-values are approximately uniform (Supplementary Fig. 3; median p-value 0.493), confirming proper calibration on an independent dataset.

---

**Supplementary Fig. 3.**
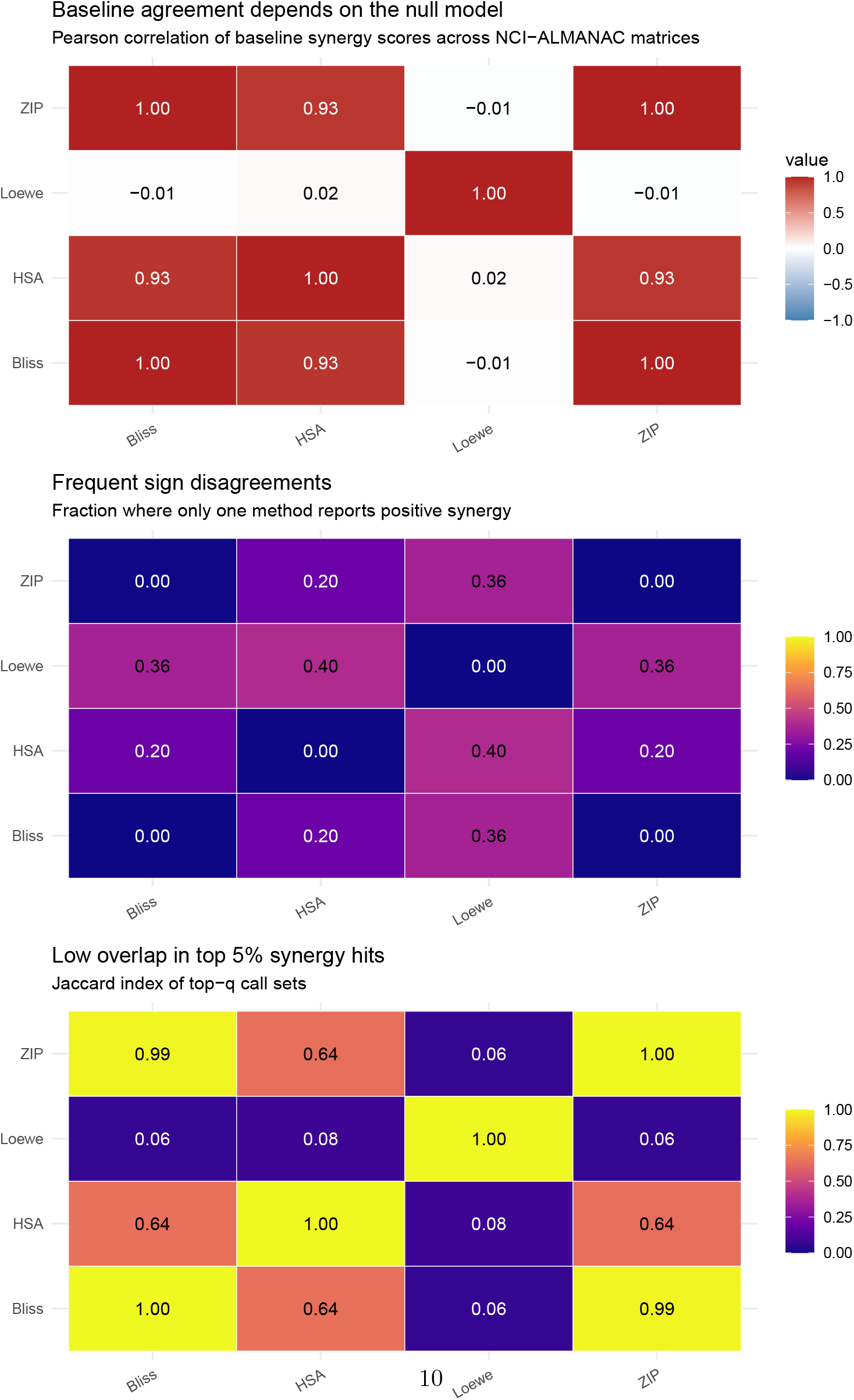
Baseline synergy score disagreement on NCI-ALMANAC. (*n* = 1,918 matrices with valid scores for all four methods). Top: Pearson correlations of matrix-level summary scores. Middle: fraction of matrices where only one method reports positive synergy. Bottom: Jaccard overlap of top 5% synergy calls. The disagreement pattern replicates and amplifies the Drug-CombDB results (main text Fig. 1): Bliss and ZIP are near-identical (*r >* 0.99), while Loewe disagrees with all other methods (*r ≈* 0, Jaccard overlap 0.06), reflecting the fundamentally different dose-equivalence assumption.

**Supplementary Fig. 4.**
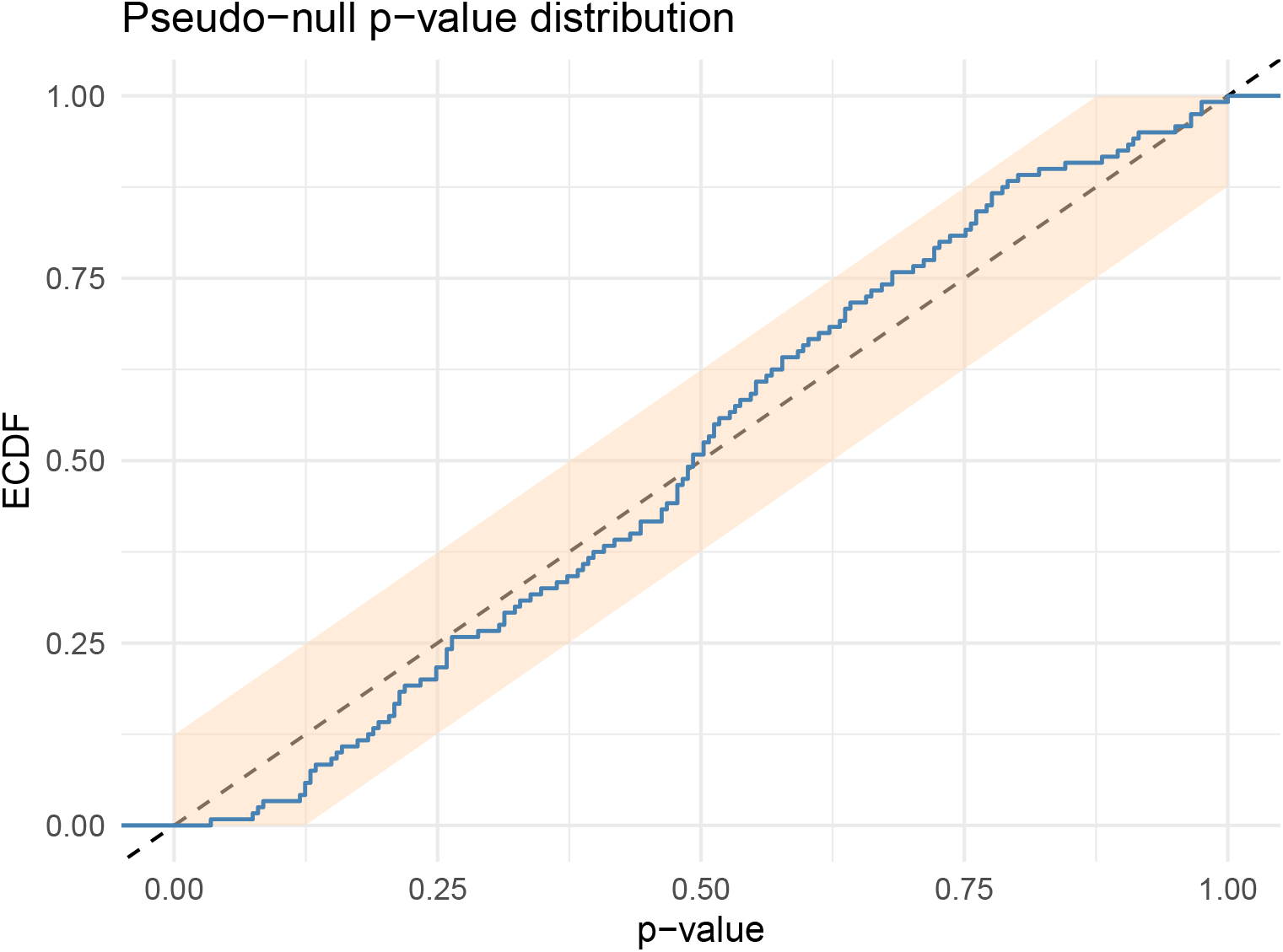
Pseudo-null p-value distribution on NCI-ALMANAC. (*n* = 120 matrices; *B* = 200). P-values from the df-corrected wild bootstrap when data are generated under the fitted additive null by sign-flipping corrected residuals, so that no true interaction is present by construction. The dashed diagonal is the Uniform(0,1) reference; the shaded region is a 95% Dvoretzky–Kiefer–Wolfowitz band. The close agreement with the uniform diagonal (median p-value 0.493) confirms that SIR’s calibration, demonstrated on DrugCombDB in the main text (Fig. 3), generalizes to an independent dataset with different experimental protocols and dose-response characteristics.

